## Supplemental figures for "CRISPR/Cas9-Mediated Excision of ALS/FTD-Causing Hexanucleotide Repeat Expansion in *C9ORF72* rescues major disease mechanisms *in vivo* and *in vitro*"

**Supplementary Figure 1**

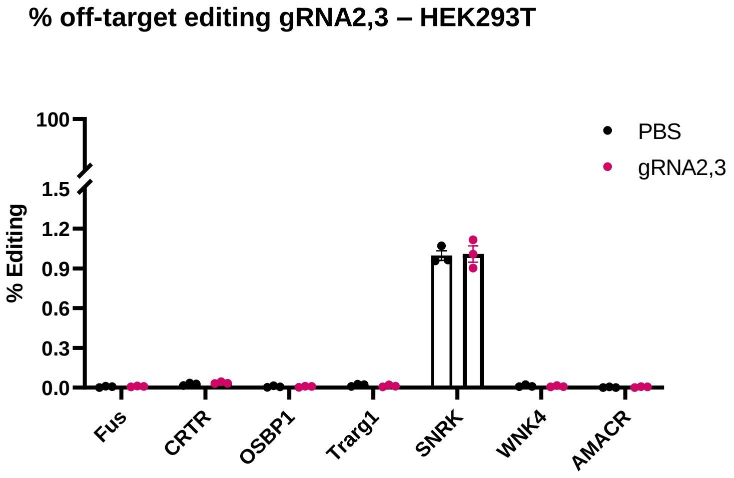


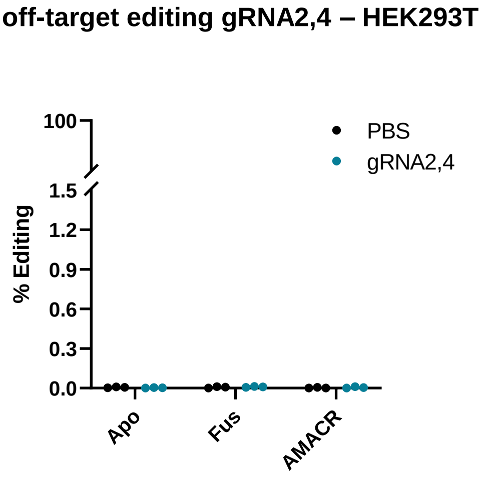


**Supplementary Figure 1.** CRISPR Amplicon Sequencing results after gRNA2,3 and gRNA2,4 editing in HEK293T cells of coding sequences that had 3 mismatches with gRNA 2, 3 or 4. NGS data was analyzed using CRISPResso2, $n$ = 3, no significant differences between treated and untreated cells.

**Supplementary Figure 2**

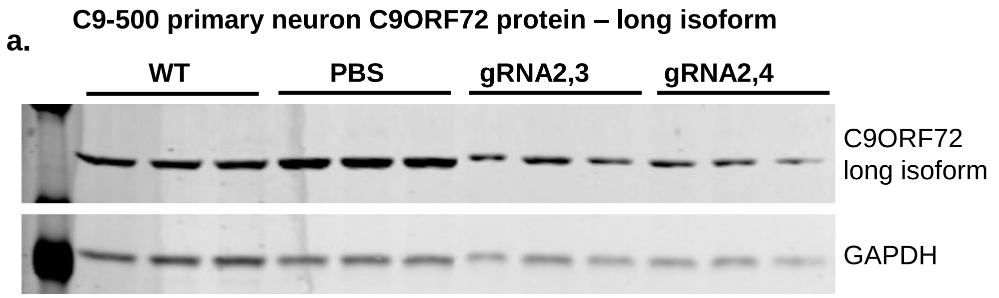


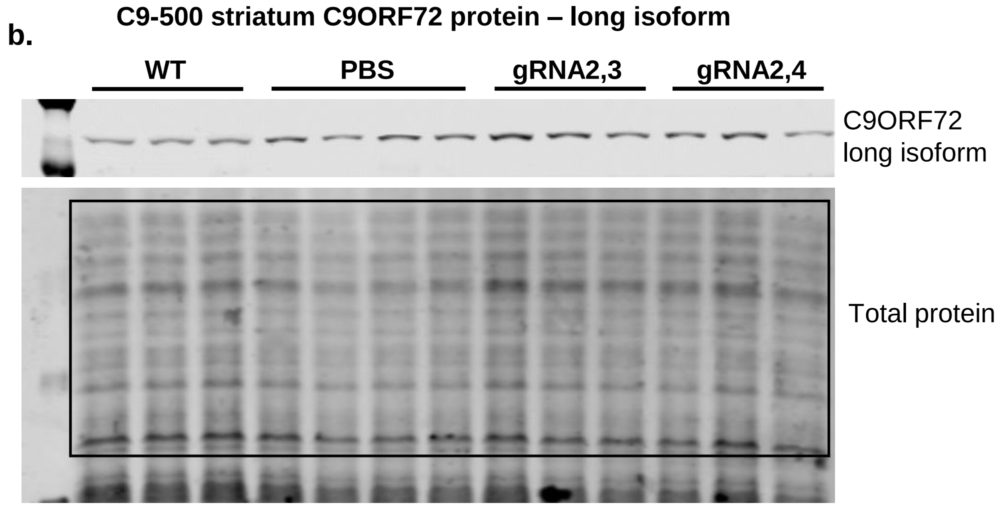


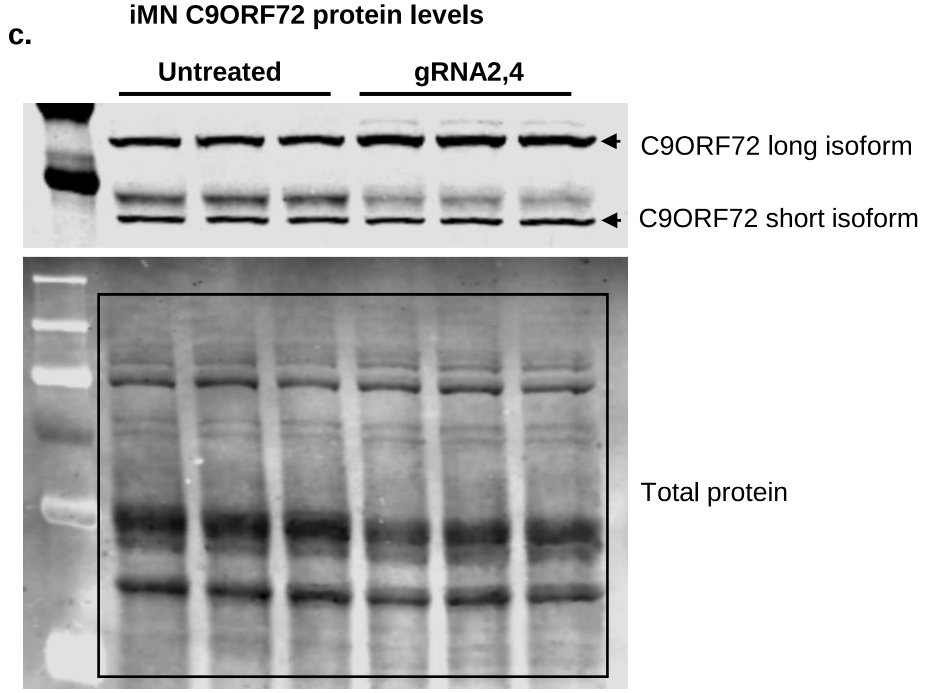


**Supplementary Figure 2.** Original blots of Western blot. **a.** Original blots of Figure 2g. **b.** Original blots of Figure 3h. **c.** Original blots of Figure 4i.

### Supplementary Figure 3

**RNA foci – non-transgenic primary neurons**

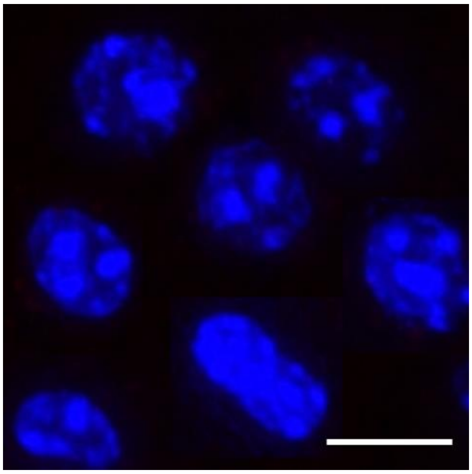

**RNA foci – non-transgenic striatum**

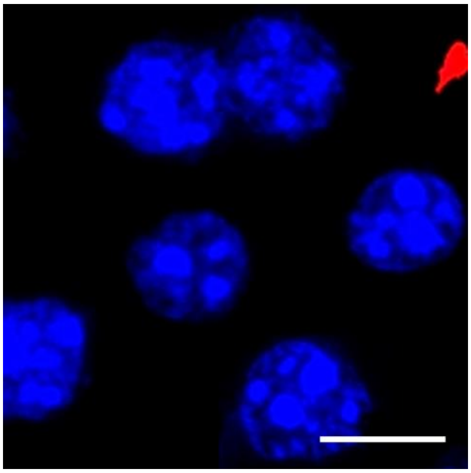

**Supplementary Figure 3.** Representative images of non-transgenic primary neurons and non-transgenic striatum stained for sense RNA foci using (FISH). Stainings for non-transgenic primary neurons and striatums were performed at the same experiment as Fig. 2d. C9-500 primary neurons and Fig. 3e. C9-500 striatums. to No sense RNA foci were detected in non-transgenic samples. Scale bars represent 10μm.
